# On the similarities of representations in artificial and brain neural networks for speech recognition

**DOI:** 10.1101/2022.06.27.497678

**Authors:** Cai Wingfield, Chao Zhang, Barry Devereux, Elisabeth Fonteneau, Andrew Thwaites, Xunying Liu, Phil Woodland, William Marslen-Wilson, Li Su

**Author notes:** These authors contributed equally to this work (Cai Wingfield, Chao Zhang).

## Abstract

How the human brain supports speech comprehension is an important question in neuroscience. Studying the neurocomputational mechanisms underlying human language is not only critical to understand and develop treatments for many human conditions that impair language and communication but also to inform artificial systems that aim to automatically process and identify natural speech. In recent years, intelligent machines powered by deep learning have achieved near human level of performance in speech recognition. The fields of artificial intelligence and cognitive neuroscience have finally reached a similar phenotypical level despite of their huge differences in implementation, and so deep learning models can—in principle—serve as candidates for mechanistic models of the human auditory system. Utilizing high-performance automatic speech recognition systems, and advanced noninvasive human neuroimaging technology such as magnetoencephalography and multivariate pattern-information analysis, the current study aimed to relate machine-learned representations of speech to recorded human brain representations of the same speech. In one direction, we found a quasi-hierarchical functional organisation in human auditory cortex qualitatively matched with the hidden layers of deep neural networks trained in an automatic speech recognizer. In the reverse direction, we modified the hidden layer organization of the artificial neural network based on neural activation patterns in human brains. The result was a substantial improvement in word recognition accuracy and learned speech representations. We have demonstrated that artificial and brain neural networks can be mutually informative in the domain of speech recognition.

**Author summary:** The human capacity to recognize individual words from the sound of speech is a cornerstone of our ability to communicate with one another, yet the processes and representations underlying it remain largely unknown. Software systems for automatic speech-to-text provide a plausible model for how speech recognition can be performed. In this study, we used an automatic speech recogniser model to probe recordings from the brains of participants who listened to speech. We found that the parts of the dynamic, evolving representations inside the machine system were a good fit for representations found in the brain recordings, both showing similar hierarchical organisations. Then, we observed where the machine’s representations diverged from the brain’s, and made experimental adjustments to the automatic recognizer’s design so that its representations might better fit the brain’s. In so doing, we substantially improved the recognizer’s ability to accurately identify words.

## Introduction

Speech comprehension—the ability to accurately identify words and meaning in a continuous auditory stream—is a cornerstone of the human communicative faculty. Nonetheless, there is still limited understanding of the neurocomputational representations and processes in the human brain which underpin it. One way to approach this question is in reverse: to find artificial systems which can accomplish the task, and use them to model and probe the brain’s solution. In the domain of engineering, automatic speech recognition (ASR) systems are designed to identify words from recorded speech audio. In this way, ASR systems provide a computationally explicit account of how speech recognition *can* be achieved, so correspondences between the human and machine systems are of particular interest; specifically, the question of whether the learned representations in an ASR can be linked to those found in human brains. Modern advances in high-resolution neuroimaging and multivariate pattern-information analysis have made this investigation feasible.

In the present research, we took a bidirectional approach, relating machine-learned representations of speech to recorded brain representations of the same speech. First, we used the representations learned by an ASR system with deep neural network (DNN) acoustic models [24] to probe the representations of heard speech in the brains of human participants undergoing continuous brain imaging. This provided a mechanistic evidence of speech responses in human auditory cortex. Then, in the opposite direction, we used the architectural patterns of neural activation we found in the brains to refine the DNN architecture and demonstrated that this improves ASR performance. This bidirectional approach was made possible by recently developed multivariate pattern analysis methods capable of comparing learned speech representations in living brain tissue and computational models.

### A computational model of speech recognition

ASR encompasses a family of computationally specified processes which perform the task of converting recorded speech sounds to the underlying word identities. Modern ASR systems employing DNN acoustic and language models now approach human levels of word recognition accuracy on specific tasks. For instance, regarding English, the word error rate (WER) of transcribing careful reading speech with no background noise can be lower than 2% [35, 48], and the WER of transcribing spontaneous conversational telephone speech can be lower than 6% [52, 66].

For the present study, our ASR system was constructed based on a set of hidden Markov models (HMMs). For each, a designated context-dependent phonetic unit handled the transitions between the hidden states. A DNN model was used to provide the observation probability of a speech feature vector given each HMM state. This framework is often called a “hybrid system” in the ASR literature [6, 24].The Hidden Markov Model Toolkit (HTK: [67, 70]), among the most widely used ASR software, was used to train the DNN-HMMs and construct the overall ASR pipeline of audio to text. A version of this model comprised a key part of the first-place winner of the multi-genre broadcast (MGB) challenge of the IEEE Automatic Speech Recognition and Understanding Workshop 2015 [4, 64]. In this paper, all ASR systems were built in HTK using 200 hours of training data from the MGB challenge. We designed the experimental setup carefully to use only British English speech and reduce the channel difference caused by different recording devices.

Of particular importance for the present study is the inclusion of a low-dimensional *bottleneck* layer in the DNN structure of our initial model. Each of the first five hidden layers contains 1000 nodes, while the sixth hidden layer has just 26 nodes. Since the DNN layers are feedforward and fully connected, each node in each layer is connected only with the nodes from its immediately preceding layer, and as such the acoustic feature representations of the input speech are forced to pass through each layer in turn to derive the final output probabilities of the context-dependent phonetic units. The bottleneck layer representations are highly compressed and discriminative, and are therefore widely used as an alternative type of input features to acoustic models in ASR literature [21, 60, 64]. In addition, the inclusion of this bottleneck layer greatly reduces the number of DNN parameters without significantly diminishing the accuracy of word recognition [64], since it can prevent the model from over-fitting to the training data [5]. Thus, the bottleneck layer representation provides a learned, low-dimensional representation of speech which is both parsimonious and sufficient for high-performance speech recognition. This is especially interesting for the present study, given the inherently low-dimensional parametrisation of speech that is given by articulatory features, which are a candidate characterisation of responses to speech in human auditory cortex.

### Speech responses in human auditory cortex

Recent electrocorticography (ECoG; [9, 18, 37, 38, 43, 44]) and functional magnetic resonance imaging (fMRI; [1, 13]) studies in humans show differential responses to speech sounds exhibiting different articulatory features in superior temporal speech areas. Heschl’s gyrus (HG) and surrounding areas of the bilateral superior temporal cortices (STC) have also shown selective sensitivity to perceptual features of speech sounds earlier in the recognition process [8, 40, 50, 56, 57]. Building on our previous work investigating phonetic feature sensitivity in human auditory cortex [63], we focus our present analysis within language-related brain regions: STC and HG.

The neuroimaging data used in this study comes from electroencephalography and magnetoencephalography (EMEG) recordings of participants listening to spoken words in a magnetoencephalography (MEG) brain scanner. High-resolution magnetic resonance imaging (MRI) was acquired using a 3T MRI scanner for better source localization. As in our previous studies [19, 56, 63], the data (EMEG and MRI) has been combined to generate a source-space reconstruction of the electrophysiological activity which gave rise to the measurements at the electroencephalography (EEG) and MEG sensors. Using standard minimum-norm estimation (MNE) procedures guided by anatomical constraints from structural MRIs of the participants [20, 23], sources were localised to a cortical mesh at the grey-matter–white-matter boundary. Working with source-space activity allows us to retain the high temporal resolution of EMEG, while gaining access to resolved spatial pattern information. It also provides the opportunity to restrict the analysis to specific regions of interest on the cortex, where an effect of interest is most likely to be found.

### Multivariate methods for modelling dynamic brain states

Recent developments in multivariate neuroimaging pattern analysis methods have made it possible to probe the representational content of recorded brain activity patterns. Among these, representational similarity analysis (RSA: [33]) provides a flexible approach which is well suited to complex computational models of rich stimulus sets. The fundamental principle of our RSA procedures was the computation of the similarity structures of the brain’s response to experimental stimuli, and comparing the similarity structures with those derived from computational models. In a typical RSA study, this similarity structure is captured in a representational dissimilarity matrix (RDM), a square symmetric matrix whose rows and columns are indexed by the experimental stimuli, and whose entries give values for the dissimilarity of two conditions, as given by their correlation distance in the response space.

A key strength of RSA is that RDMs abstract away from the specific implementation of the DNN model or measured neural response, allowing direct comparisons between artificial and human speech recognition systems; the so-called “dissimilarity trick” [32]. The comparison between RDMs computed from the ASR model and RDMs from human brains take the form of a Spearman’s rank correlation *ρ* between the two [46].

RSA has been extended using the fMRI searchlight-mapping framework [31, 46] so that representations can be mapped through image volumes. Subsequently, searchlight RSA has been further extended into the temporal dimension afforded by EMEG data: spatiotemporal searchlight RSA (ssRSA: [55, 56]). Here, as in other studies using computational cognitive models (e.g. [27, 36]), ssRSA facilitates the comparison to a machine representation of the stimulus space which may otherwise be incommensurable with a distributed brain response.

### Outline

In the machine-to-human direction, using ssRSA and the ASR system as a reference, we found that the early layers of the DNN corresponded to early neural activation in primary auditory cortex, *i*.*e*. bilateral Heschl’s gyrus, while the later layers of the DNN corresponded to late activation in higher level auditory brain regions surrounding the primary sensory cortex. This finding reveals that the neural network located within HG is likely to have a similar functional role as early layers of the DNN model, extracting basic acoustic features. The neurocomputational function of superior temporal gyrus regions is akin to later layers of the DNN, computing complex auditory features such as articulation and phonemic information.

In the reverse human-to-machine direction, using the pattern of results in the brain-image analysis, we improved the architecture of the DNN. The spatial extent of neural activation explained by the hidden-layer representations progressively reduced for higher layers, before expanding again for the bottleneck layer. This pattern, which mirrored the structure of the DNN itself, and (assuming an efficient and parsimonious processing stream in the brain) suggests that some pre-bottleneck layers might be superfluous in preparing the low-dimensional bottleneck compression. We restructured the DNN model with the bottleneck layer moved to more closely resemble the pattern of activation observed in the brain, hypothesising that this would lead to a better transformation. With this simple, brain-inspired modification, we significantly improved the performance of the ASR system. It is notable that similar DNN structures have been developed independently elsewhere in order to optimise the low-dimensional speech feature representations from the DNN bottleneck layer. However, “reverse-engineer” human learning systems implemented in brain tissue in such a bidirectional fashion provides a complementary approach in developing and refining DNN learning algorithms.

## Results

### DNN bottleneck layer activations are organized by articulatory features

A DNN acoustic model was trained to classify each input frame into one of the triphone units at each time step. The DNN had five 1000-node hidden layers followed by a single 26-node bottleneck (BN) layer, and is therefore denoted as DNN-BN_7_ since the bottleneck layer is the seventh layer (L7). We used it as the acoustic model of our DNN-HMM ASR system to estimate the triphone unit likelihoods corresponding to each frame. The log-Mel filter bank (FBK) acoustic features were used throughout the paper, which were extracted with a 25 ms duration and 10 ms frame shift. The first order differentials of the FBK features were also included to extend the acoustic feature vectors. Nine consecutive frames of the acoustic features centred at the current time step were stacked to form the DNN input vector, which covers a total range of 125 ms of the speech signal. More details about the DNN acoustic model can be found in the Methods Section.

Our hypothesis was that the representations in auditory cortex would be organised according to phones and features [63]. To investigate how the assignment of phonetic and featural labels to each segment of the stimuli could explain hidden-layer representations in DNN-BN_7_, we computed Davies–Bouldin clustering indices for representational spaces at each layer. Davies– Bouldin clustering indices give an indication of the degree to which a layer’s response to each segment of audio form clusters which correspond to a set of category labels. This in turn serves as an indication of how suitably phonetic and feature labels might be assigned to hidden-layer representations.

Davies–Bouldin indices for each layer and categorisation scheme are shown in Fig 1A. Of particular interest is the improvement of feature-based clustering in bottleneck layer L7 of DNN-BN_7_, which shows that it is, in some sense, reconstructing the featural *articulatory* dimensions of the the speaker. That is, though this was not included in the teaching signal, when forced to parsimoniously pass comprehension-relevant information through the bottleneck, DNN-BN_7_ finds a representation of the input space which maps well onto the constraints on speech sounds inherent in the mechanics of the speaker. L7 showed the best clustering indices out of all layers for manner and place features and phone labels, and the second-best for frontness features. For closeness alone, L7 was not the best, but was still better than its adjacent layer L6. The general trend was that clustering improved for successively higher layers. Layers prior to the bottleneck tended to have larger clustering indices, indicating that their activations were not as well accounted for by phonetic or featural descriptions.

**Figure 1:**
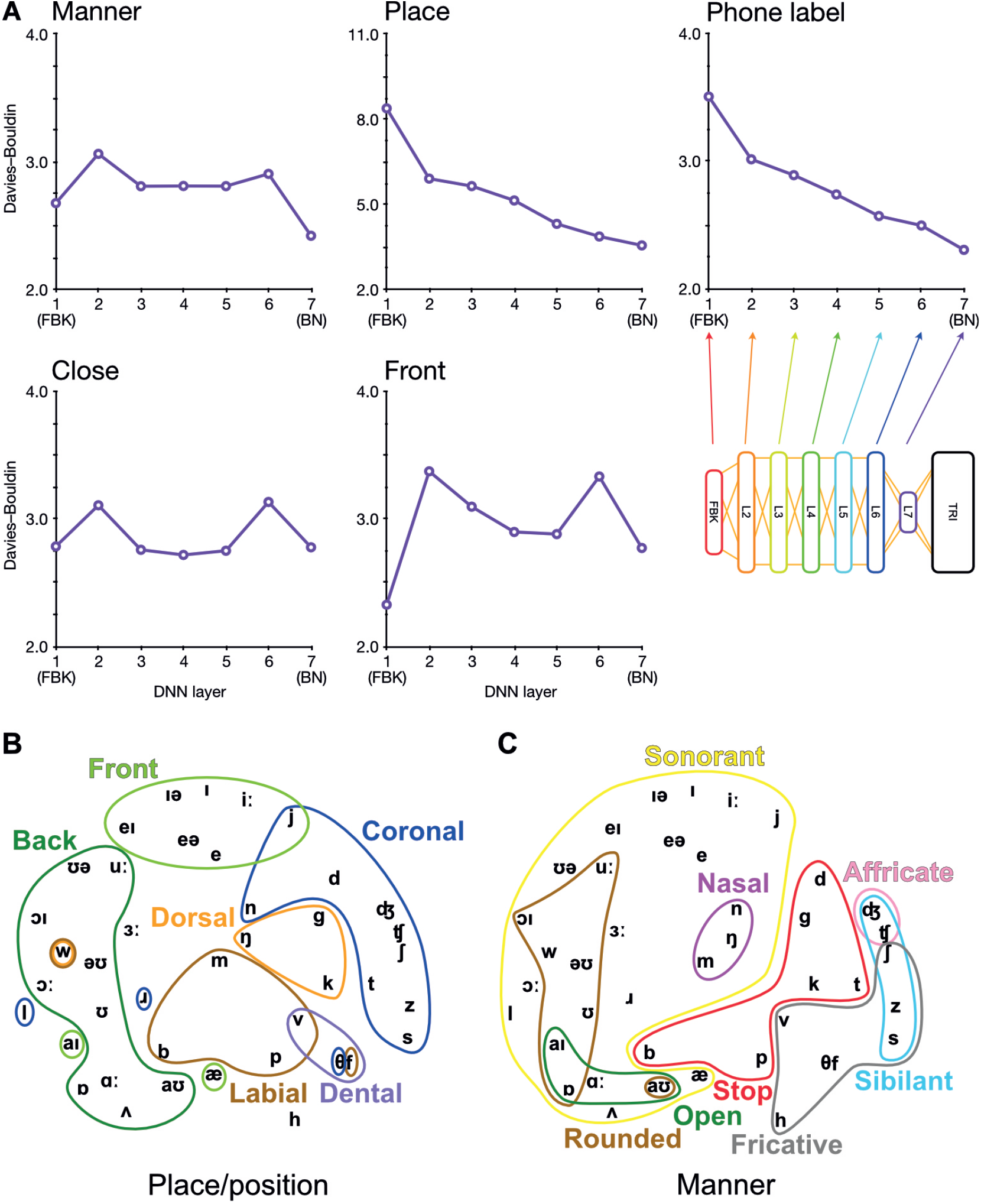
Arrangement of phonetic space represented in DNN-BN_7_. **(A)** Davies–Bouldin clustering indices for hidden-layer representations. Each plot shows the Davies–Bouldin clustering index for the average hidden-layer representation for each phonetic segment of each stimulus. Lower values indicate better clustering. Indices were computed by labelling each segment by its phonetic label (top right panel), or by place, manner, frontness or closeness features (other panels). Colored shapes on the DNN-layer axis indicate the placement of the bottleneck layer for each System. **(B) Average activation of phones for L7** Sammon nonlinear multidimensional scaling (MDS) of average pattern of activation over phones, annotated with features describing place and position of articulation. **(C)** The same MDS arrangement annotated with features describing manner of articulation.

To further illustrate and visualise the representational space for L7, we used the phonetic partitioning of our stimuli provided by HTK, and averaged the activation across hidden nodes in L7 for each window of our 400 stimulus words which was eventually labeled with each phone. This gave us an average L7 response vector for each phone. We visualised this response space using Sammon nonlinear multidimensional scaling (MDS; [51]). Place/position features are highlighted in Fig 1B, and manner features are highlighted in Fig 1C.

To be clear, the presence of these feature clusters does not imply that there are individual nodes in L7 which track specific articulatory features. However, using the reasoning of RSA, we can see that articulatory features are descriptive of the overall arrangement of phones in the L7 response space. This ability to characterize and model an overall pattern ensemble in a way abstracted from the specific response format and distributed neural representations is one of the strengths of the RSA technique.

### Hidden-layer representations differentially explain early human auditory cortex representations through space and time

We used the dynamic representations from each layer of DNN-BN_7_ to model spatiotemoral representations in the auditory cortices of human participants in an EMEG study by applying ssRSA. Areas of auditory cortex (Fig 2A) were defined using the Desikan–Killiany Atlas (STC and HG).

**Figure 2:**
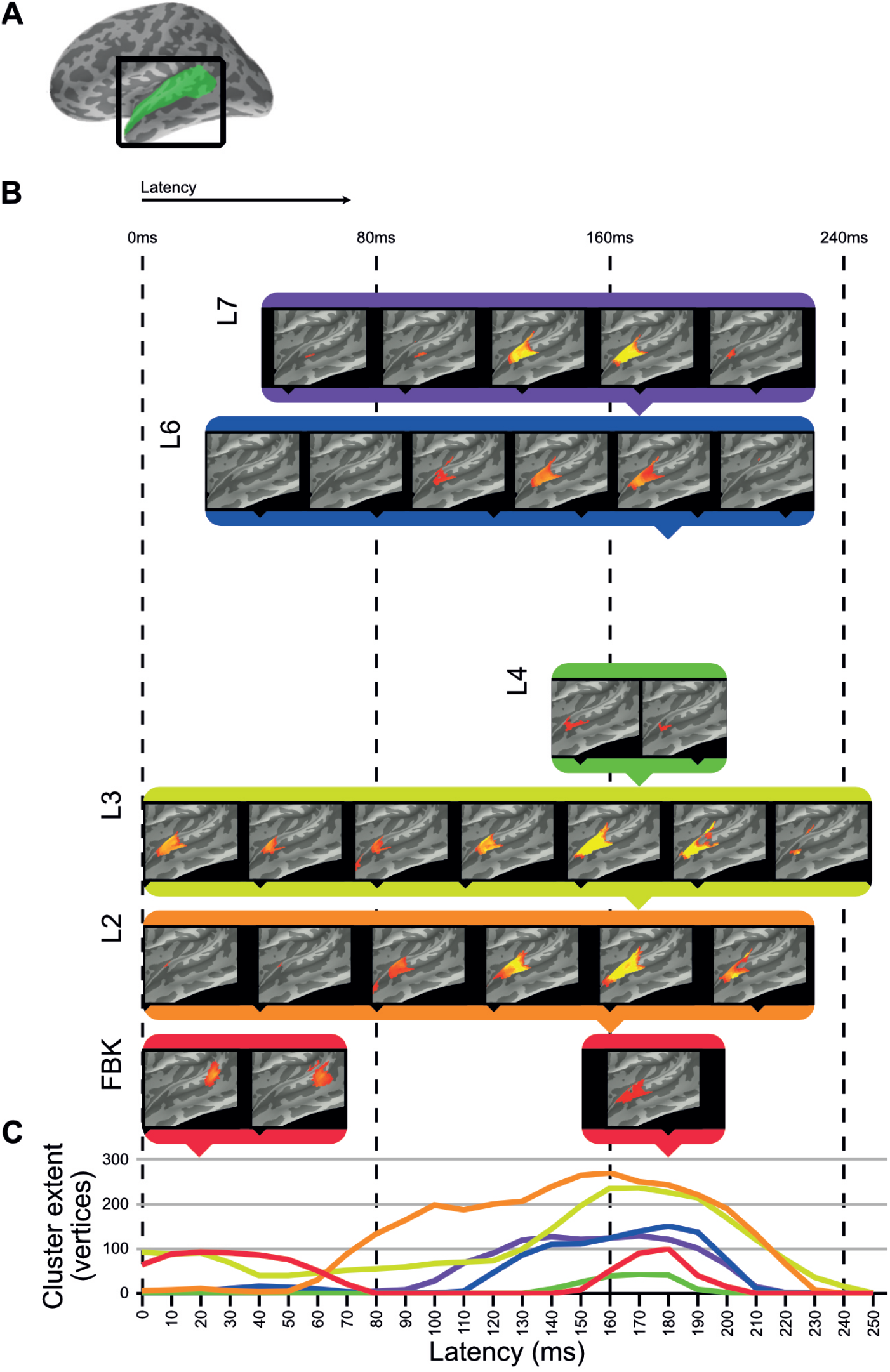
Clusters of significant fit of hidden-layer models to left-hemisphere EMEG data. **(A)** Location of region of interest mask for auditory cortex. **(B)** Maps describing fit of DNN layer models to EMEG data. Latency represents the time taken for the brain to exhibit neural representations that fit the DNN model prediction. All maps thresholded at *p <* 0.01 (corrected). **(C)** Line graphs showing the time-courses of cluster extents for each layer which showed significant fit.

Fig 2 shows the left hemisphere results of this analysis. The brain maps in Fig 2B show thresholdfree-cluster-enhanced *t*-maps [54] computed from the model RDMs of each hidden layer, thresholded at *p <* 0.01. Model RDMs computed from all DNN layers except L5 showed significant fit in left STC and HG. Input layer FBK peaked early in left posterior STC at 0–70 ms, and later in left anterior STC and HG at 140–210 ms. Hidden-layer models L2–L4 and L6–L7 peaked later than FBK, achieving maximum cluster size at approximately 170 ms. Layers L5 and TRI showed no significant fit in the regions of interest. Overall, significant cluster size increased between layers FBK–L3, diminished for L4 and L5, and re-emerged for L6 and L7.

The line graphs in Fig 2C show the time-courses of each layer as they attain their maximum cluster extent. In general, there appeared to be two distinct peaks across the superior temporal region: an early peak in left posterior STC for the DNN input layer FBK, and another late peak in left anterior STC for DNN layers L1–L4 and L6–L7, throughout the whole epoch, but attaining a maximum cluster size at approx 170 ms. Details of timings for each layer are shown in S2 Table. Right hemisphere results are included in S3 Fig.

### Repositioning the DNN bottleneck layer to match human brain improves ASR performance and featural organization

The overall minimal spatiotemporal clusters for L5 of DNN-BN_7_ suggested that while early layers (L2–L3) were performing analogous transformations to early auditory cortex, and that the bottleneck (L7) was representing speech audio with a similarly parsimonious basis as left auditory cortex, there was a divergence of representation at intermediate layers (L4–L6). With the supposition that the arrangement of auditory cortex would be adapted specifically to speech processing, we hypothesised that by moving the bottleneck layer into the positions occupied by divergent layers in DNN-BN_7_, the network might learn representations that closer resemble those of human cortex, and thus improve the performance of the model. To this end, we built and studied another DNN model, DNN-BN_5_, which has the same number of parameters as DNN-BN_7_ but has the bottleneck layer moved from L7 to L5 (see Fig 3C. For purposes of comparison, and following the same naming convention, we expanded our investigation with another two DNN models, DNN-BN_4_ and DNN-BN_6_ were also built for DNNs whose bottleneck layers are L4 and L6 respectively. In all models the number of parameters was kept to 5.0 million, matching the 4.9 million parameters of DNN-BN_7_.

**Figure 3:**
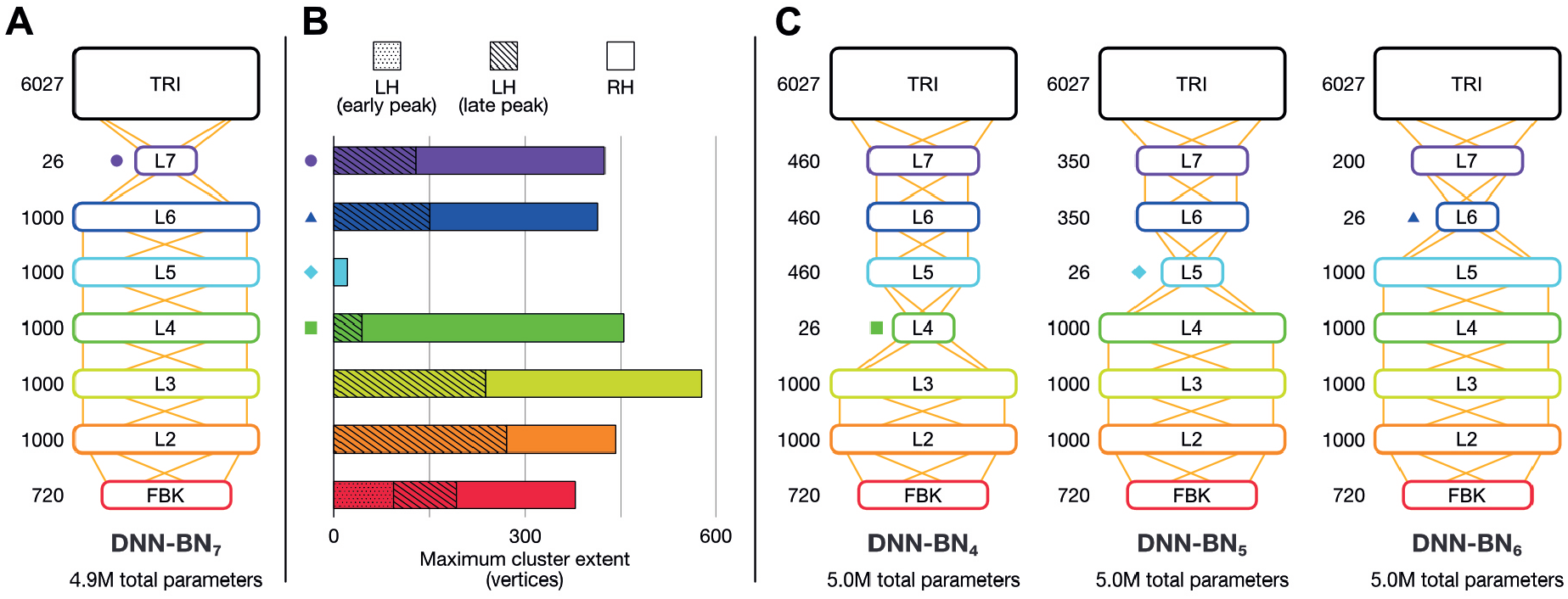
Brain-informed DNN design refinement. **(A) Original DNN-BN_7_ design.** Numbers beside layers indicate number of nodes. **(B)** Degree of fit with EMEG brain representations. Shapes here and other panels indicate bottleneck positions for DNN-BN_4–7_ **(C)** Candidates for adjusted DNN design: DNN-BN_4_ (bottleneck at L4), DNN-BN_5_ (bottleneck at L5) and DNN-BN_6_ (bottleneck at L6).

As shown in Table 1 and Fig 4C, adjusting the design of the DNN structure to better fit with the representations exhibited in the human subjects led to improved DNN performance in terms of WER in (DNN-BN_6_). The MGB Dev set contains sufficient testing samples with diversified speaker and genre variations. The 1.0% absolute WER reduction (relatively 3.3%) obtained by comparing DNN-BN_7_ with DNN-BN_5_ is substantial [4, 64]. Regarding the stimulus set, the changes of WERs are consistent with those on the MGB Dev set.

**Table 1:** The performance of DNN-HMM systems with different bottleneck layer positions. The WERs (the lower the better) were given on both the MGB challenge official development subset (MGB Dev), which is a general purpose large vocabulary continuous speech recognition testing set, as well as the 400 isolated words used as the stimuli in our listening experiments to derive the RDM (Stimuli). The MGB Dev WERs are reliable indicators for the general performance of the systems in realistic ASR tasks. The Stimuli WERs are the most direct indicators of the model performance on the data used in our brain-machine comparison experiments. The classification accuracy values (the higher the better) were obtained by classifying each frame into one of the 6,027 triphonetic DNN output units were obtained on both the training and held-out validation (HV) sets. For fair comparisons, DNN structures of all systems were constrained to have the same amount of model parameters (about 5M for each model, as shown in Figure 3). Accuracy can be considered as an auxiliary performance metric, which indicates that DNN-BN_6_ suffered more from over-fitting compared to DNN-BN_5_, since DNN-BN_6_ is better in the training accuracy but not in the HV accuracy.

| System | Bottleneck layer | Accuracy% |  | WER% |  |
| --- | --- | --- | --- | --- | --- |
|  |  | Train | HV | MGB Dev | Stimuli |
| DNN-BN<sub>7</sub> | L7 | 44.0 | 41.5 | 33.3 | 6.5 |
| DNN-BN<sub>6</sub> | L6 | 44.6 | 42.3 | 32.4 | 6.3 |
| DNN-BN<sub>5</sub> | L5 | 44.2 | 42.3 | 32.3 | 5.8 |
| DNN-BN<sub>4</sub> | L4 | 42.6 | 41.1 | 33.5 | 7.3 |

**Figure 4:**
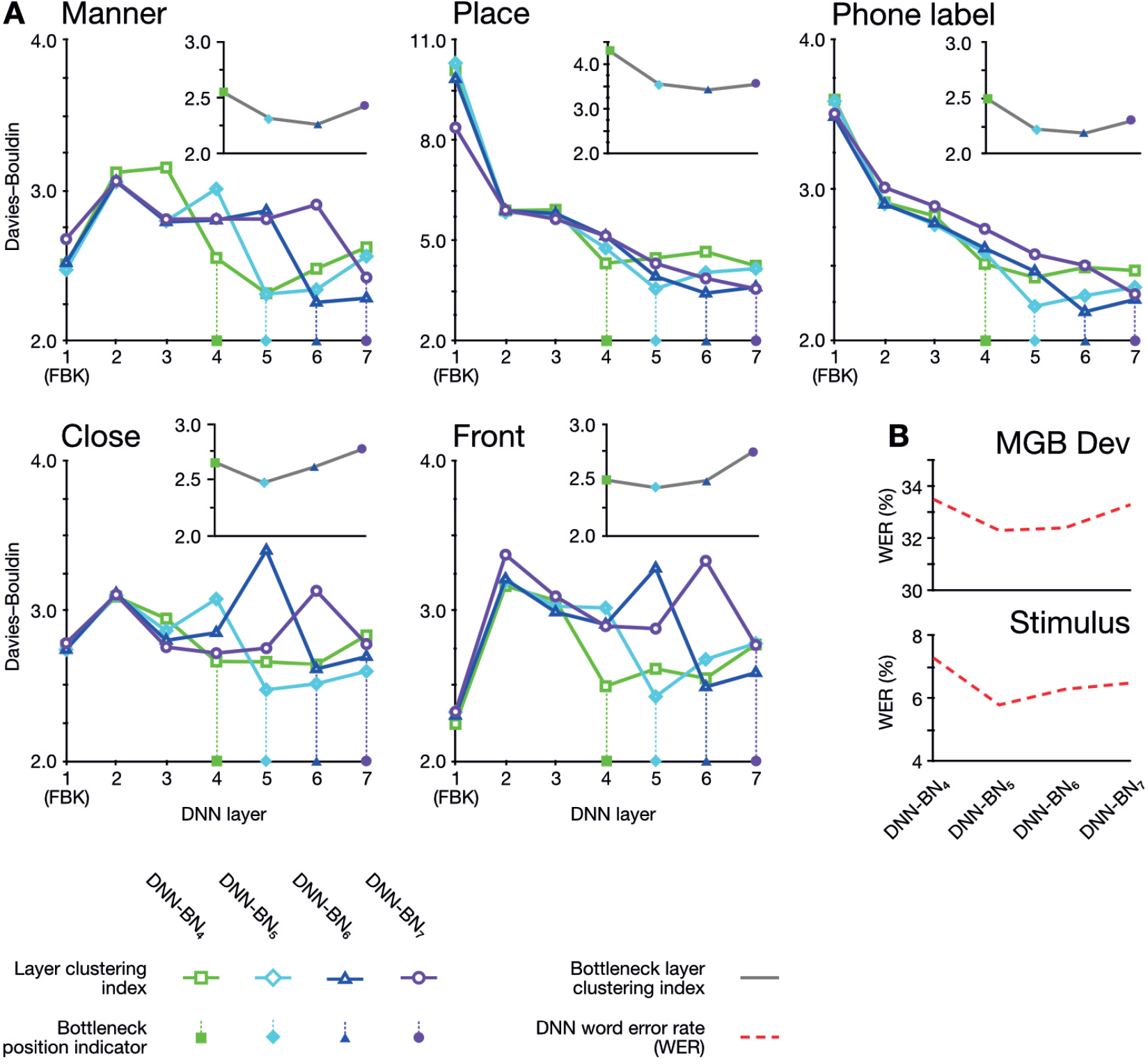
(A) Davies–Bouldin clustering indices for hidden-layer representations. Each plot shows the Davies–Bouldin clustering index for the average hidden-layer representation for each phonetic segment of each stimulus. Lower values indicate better clustering. Indices were computed by labelling each segment by its phonetic label (top right panel), or by place, manner, frontness or closeness features (other panels). Colored shapes on the DNN-layer axis indicate the placement of the bottleneck layer for each System. Inset axes show clustering indices for bottleneck-layers only. Each plot shows the clustering index for the average bottleneck-layer representation for each phonetic segment of each stimulus. Indices were computed by labeling each segment by its phonetic label (top right), or by place, manner, frontness or closeness features. Colored shapes on the DNN-layer axis indicate the placement of the bottleneck layer for each System. **(B) WERs for each DNN system**. Upper panel shows WERs on the MGB Dev set. Lower panel shows WERs for the stimuli.

What is not immediately clear, however, is whether this improvement in performance arises from a corresponding improvement in the model’s ability to extract a feature-based representation. In other words, if the bottleneck layer learns a representation akin to articulatory features, by moving the layer to improve performance does this enhance this learned representation? To answer this question, we investigated how the assignment of phonetic and featural labels to each segment of the stimuli could explain their hidden-layer representations. As before, we probed the organization of the representational space of each hidden layer according to phones and features using Davies–Bouldin clustering indices.

The clustering results exhibited two overall patterns of note. First, clustering (*i*.*e*. suitability of assignment of phonetic and featural labels to hidden layer representations) was improved on the DNNs whose design had been inspired by the human brains. Second, the optimum clustering level was often found in the bottleneck layer itself (highlighted on the graphs in Fig 4A). The clustering index at the bottleneck layers alone are separately graphed in inset panels in Fig 4A, and show that bottleneck layer clustering was also improved in DNN-BN_5_ and DNN-BN_6_.

In other words, the placement of the bottleneck layer in position 5 and 6 yielded, as predicted, the best clustering results both overall and in the bottleneck layer itself. Moving the bottleneck layer too far back (DNN-BN_4_) yielded worse clustering results generally and in the bottleneck layer—indicated by the characteristic U-shaped curves in Fig 4B.

## Discussion

We have used a DNN-based ASR system and spatiotemporal imaging data of human auditory cortex in a mutually informative study. In the machine-to-human direction, we have used a computational model of speech processing to examine representations of speech throughout space and time in human auditory cortex measured as source-localised EMEG data. In so doing, we have produced a functional map in human subjects for each part of the multi-stage computational model. We were able to relate dynamic states in the operating machine speech recognizer to dynamic brain states in human participants by using ssRSA, extended to account for a dynamically changing model. In a complementary analysis, we have improved the performance of the DNN-based ASR model by adapting the layered network architecture inspired by the staged neural activation patterns observed in human auditory cortex.

### Locations of spatiotemporal clusters

The input layer FBK representing purely acoustic information (*i*.*e*. not a learned or task-relevant representation) showed a later and smaller effect (cluster in human posterior STC) than that of higher layers L2 and L3. The strongest peak for FBK was early, and the later peak appears to be a weaker version of those for higher hidden-layer models. The late peak for FBK indicates that there is some involvement of both low-level acoustic features and higher-level phonetic information in the later neural processes at around 170 ms. However, since there is an intrinsic correlation between acoustic information and phonetic information, it is hard to completely dissociate them. Another explanation for the mixture of high and low levels of speech representations in a single brain region at the same time is the existence of feedback connections in human perceptual systems. (However, the ASR systems used in this paper can achieve high degree of accuracy without the top-down feedback loop from higher to lower hidden layers.) It should be noted that while the FBK, L2 and L4 clusters all register as significant at a latency of 0 ms, timings correspond to a 25 ms window of EMEG data being matched against model state computed for the central 25 ms of 125 ms windows of audio, so only approximates the actual latency.

Moving up to hidden layers L2 and L3, we saw later clusters which fit the brain data more strongly than FBK in the left hemisphere. All hidden layers including L2 and L3 activate according to learned parameters. Progressively higher layers L4 and L5 fit with smaller clusters in human STC, with L5 showing no significant vertices at any time point (*p >* 0.01) in the left hemisphere but a very small cluster in the right hemisphere. However, the highest hidden layers L6 and L7 once again showed string fit with activations in left anterior STC.

Of particular interest is this re-emergence of fit in anterior STC to the representations in the bottleneck layer L7. In this layer of the DNN, the 1000-node representation of L6 is substantially constrained by the reduced size of the 26-node L7. In particular, the fact that ASR accuracy is not greatly reduced by the inclusion of this bottleneck layer indicates that, for the machine solution, 26 nodes provide sufficient degrees of freedom to describe a phonetic space for purposes of word recognition. This, in conjunction with the re-emergence of fit for L7 to STC representations makes the representations of this layer of particular interest. The hidden layers in the DNN learn to sequentially transform acoustic information into phonetic probabilities in a way which generalises across speakers and background acoustic conditions. There is no guarantee that the features the DNN learns to identify for recognition are comparable to those learned by the brain, so the fact that significant matches in the RDMs were found between machine and human solutions of the same problem is worthy of further consideration.

### Brain-informed ASR architecture

Artificial Intelligence (AI) and machine learning have already been extensively applied in neuroscience primarily in analysing and decoding large and complex neuroimaging or cell recording data sets. Here, DNN-based ASR systems were used as a model for developing and testing hypothesis and neuroscientific theories about how human brains perform speech recognition. This type of mechanistic or generative model—where the computational model can perform the behavioral task with realistic data (in this case, spoken word recognition)—can serve as a comprehensive framework for testing claims about neurocognitive functional organization [30] Moreover, insights can flow both ways; the neuroimaging data can also guide the exploration of the model space and lead to improvements in model performance, as we have seen.

While our use of neurological data only indirectly informed the improvements to ASR architecture, the present work can be seen as an initial step toward extracting system-level designs for neuromorphic computing from human auditory systems. This goal in itself is not new (see e.g. [58]), however the key novel element of our approach is the ability to relate the machine and human solutions in complementary directions. The power of RSA, and in particular ssRSA, to relate the different forms of representations in these systems is key in this work. In summary, the methodology illustrated here paves the way for future integration of neuroscience and AI with the two fields driving each other forwards.

### Relating dynamic brain and machine states: comparing and contrasting computational models in vision and audition

There has been some recent successes in comparing machine models of perception to human neuroimaging data. This has primarily been in the domain of visual object perception (e.g. [7, 11, 12, 17, 22, 27–29, 34]), with less progress made in speech perception (though see our previous work; [56, 63]).

The visual systems of humans and other primates are highly related, both in their architecture and in accounts of the neurocomputational processes they facilitate. There is evidence of a hierarchical organisation of cortical regions in the early visual systems of human and nonhuman primates. There are also detailed accounts of process sequencing from early visual cortex through higher perceptual and semantic representation which exist for visual object perception in several primate models (e.g. [16, 34, 47, 59, 61]). This is not so the case for speech processing and audition to the same degree.

In parallel, machine models for vision have often been designed based on theories of primate cortical processing hierarchies. This extends to recent work employing deep convolutional neural networks (CNN) for visual object processing, in particular those featuring layers of convolution and pooling. Furthermore, the convolutional layers in CNNs appear to learn features resembling those in the receptive fields of early visual cortex, and higher layers’ representational spaces also match those found in higher visual cortex, and other regions in the visual object perception networks [22, 27, 62]. Importantly, this means that the internal structures of machine vision systems are potentially informative and relevant to our understanding of the neurocomputational architecture of the natural system (and vice versa), and not just whether they generate equivalent outputs (for example in object classification tasks). To date, these common features are not well established for DNNs or other type of acoustic models widely used for ASR systems.

Certain aspects of the human auditory processing system have resemblances to those in other primate models [3, 49]. However, no non-human primate supports anything like human speech communication, where intricately modulated sequences of speech sounds map onto hundreds of thousands of learned linguistic elements (words and morphemes), each with its own combination of acoustic-phonetic identifiers.

Perhaps due to this lack of neurocomputationally explicit models of spoken word recognition, the design of ASR systems has typically not been guided by existing biological models. Rather, by optimising for engineering-relevant properties such as statistical learning efficiency, they have nonetheless achieved impressive accuracy and robustness.

It is striking, therefore, that we have been able to show that the regularities that successful ASR systems encode in the mapping between speech input and word-level phonetic labelling can indeed be related to the regularities extracted by the human system. In addition, like animal visual systems have inspired the field of computer vision, we have demonstrated that human auditory cortex can improve ASR systems using ssRSA.

### Conclusion and future work

We have shown that our deep artificial neural network model of speech processing bears resemblance to patterns of activation in the human auditory cortex using the combination of ssRSA with multimodal neuroimaging data. The results also showed that the low-dimensional bottleneck layer in the DNN could learn representations that characterize articulatory features of human speech. In ASR research, although the development of systems based around the extraction of articulatory features has a long history (e.g. [15]), except for a small number of exemplars (e.g. [39, 69]), recent studies mostly rely on written-form-based word piece units [53, 65] that are not directly associated with phonetic units. Our findings imply that developing appropriate intermediate representations for articulatory features may be central to speech recognition in both human and machine solutions. In human neuroscience studies, this account is consistent with previous findings of articulatory feature representation in the human auditory cortex [13, 37, 63], but awaits further investigation and exploitation in machine solutions for speech recognition. Recently, large deep artificial neural network models pre-trained on a massive amount of unlabelled waveform features (e.g. [2, 10, 25]), have demonstrated strong generalisation abilities to ASR and many para-linguistic speech tasks [41]. It would be useful to apply our methods used in this paper to study similar types of models and tasks. This may contribute to understanding the hierarchical structures in the human auditory cortex and improve such large scale speech-based computational models.

## Materials and methods

### Deep neural networks for automatic speech recognition

We have presented four DNNs which can each be included as a component in the hybrid DNNHMM set-up of HTK. This is a widely used speech recognition set-up in both academic and industrial communities [24], whose architecture is illustrated in Fig 5. Each network comprises an input layer, six hidden layers, and an output layer, which are all fully-connected feed-forward layers.

**Figure 5:**
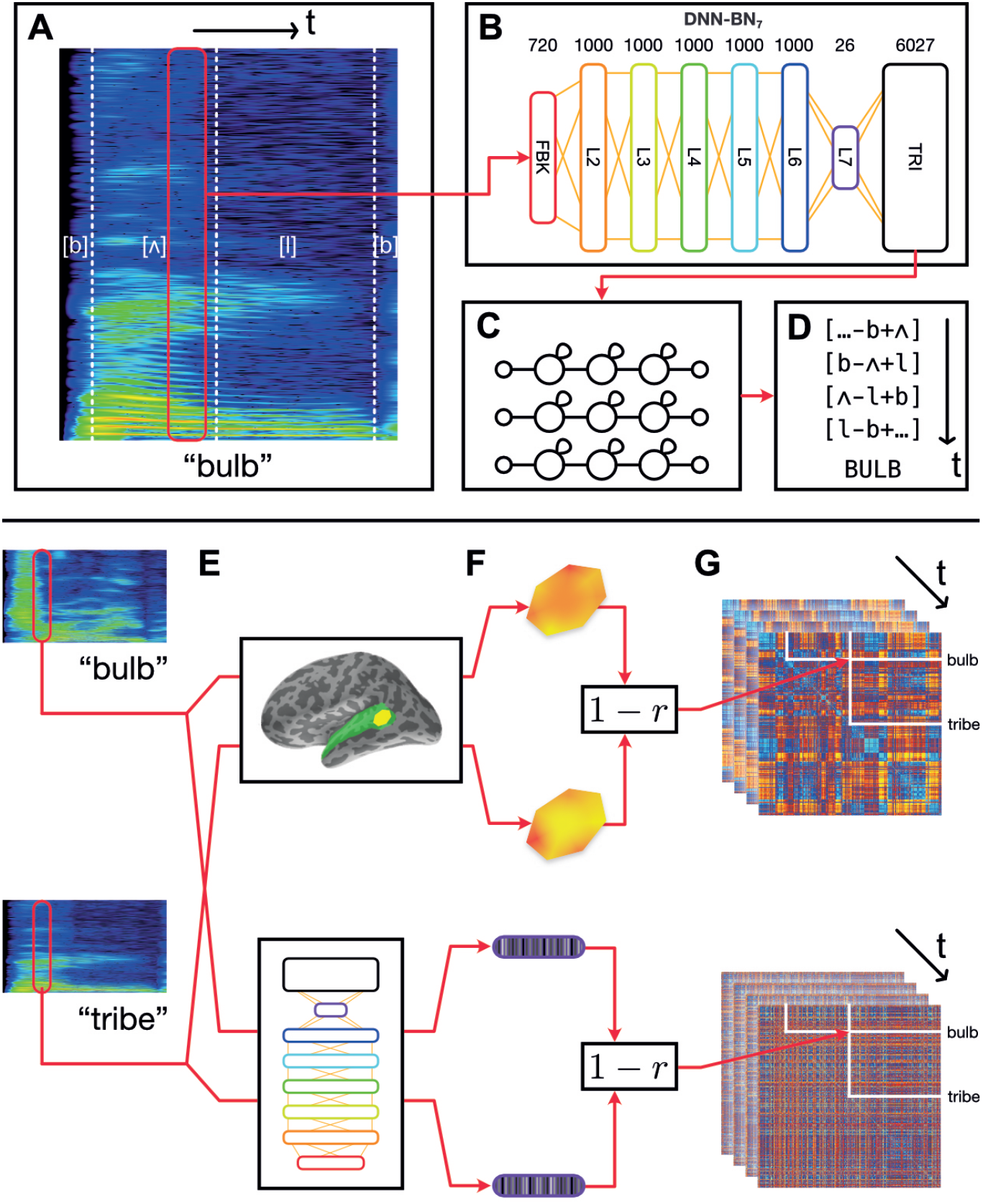
Schematic of the overall procedure. **A–D: Automatic speech recognition system HTK.** Our ASR model is a hybrid DNN–HMM system built with HTK. **(A)** An acoustic vector is built from a window of recorded speech. **(B)** This is used as an input for a DNN acoustic model which estimates posterior probabilities of triphonetic units. Numbers above the figure indicate the size of each layer. Hidden layer L7 is the bottleneck layer for DNN-BN_7_. **(C)** The triphone posteriors (TRI) are converted into log likelihoods, and used in a set of phonetic HMMs. **(D)** A decoder computes word identities from the HMM states. **E–G: Computing dynamic RDMs. (E)** A pair of stimuli is presented to each subject, and the subjects’ brain responses are recorded over time. The same stimuli are processed using HTK, and the hidden-layer activations recorded over time. **(F)** The spatiotemporal response pattern within a patch of each subject’s cortex is compared using correlation distance. The same comparison is made between hidden-layer activation vectors. **(G)** This is repeated for each pair of stimuli, and distances entered into a pairwise comparison matrix called a representational dissimilarity matrix (RDM). As both brain response and DNN response evolve over time, additional frames of the dynamic RDM are computed.

### Building DNN-HMM acoustic models for ASR

As introduced previously, the input audio stream is divided into 25 ms-long overlapping windows. Each of these windows is transformed into a 40-dimensional FBK feature vector representing a speech frame with an offset of 10 ms. When being fed into the DNN input layer, the 40-dimensional feature vectors are augmented with their first-order time derivatives (also termed as *delta features* in speech recognition literature) to form an 80-dimensional vector *o*_*t*_ for the *t*-th frame. The final DNN input feature vector, *x*_*t*_, is formed by stacking nine consecutive acoustic vectors around *t, i*.*e. x*_*t*_ = {o_*t−*4_, *o*_*t−*3_, …, *o*_*t*+4_}. Therefore, the DNN input layer (denoted as the FBK layer from Figure 1 to Figure 5) has 720 nodes and covers a 125 ms long input window starting at (10 *× t −* 50) ms and ending at (10 *× t* + 75) ms. Where this wider context window extended beyond the limits of the recording (*i*.*e*. at the beginning and end of the recording), boundary frames were duplicated to make up the nine consecutive frames.

Following the input layer FBK, there are five 1000-node hidden layers (L2–L6), a 26-node “bottleneck” layer (L7), and the output layer (TRI). All hidden nodes use a sigmoid activation function and the output layer uses a softmax activation function to estimate pseudo posterior probabilities for 6,027 output units. There are 6,026 such units corresponding to the tied triphone HMM states which are obtained by the decision tree clustering algorithm [68]. The last output unit is relevant to the non-speech HMM states. The DNN was trained on a corpus consisting of 200 hours of British English speech selected from 7 weeks of TV broadcast shows by the BBC covering all genres. Using such a training set with a reasonably large amount of realistic speech samples guarantees our DNN model to be properly trained and close to the models used in real-world speech recognition applications. The DNN model was trained to classify each of the speech frames in the training set into one of the output units based on the cross-entropy loss function.

When performing speech recognition at test-time, the posterior probabilities, *P* (*s*_*k*_ | *x*_*t*_), are converted to log-likelihoods to use as the observation density probabilities of the triphone HMM states. Specifically, the conversion is performed by

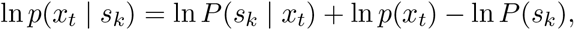

where *s*_*k*_ is a DNN output for target *k*, and *P* (*s*_*k*_) is the frequency of frames corresponding to the units associated with target *k* in the frame-to-HMM-state alignments of the training set [24].

### Recorded speech stimuli

This study used speech stimulus recordings from [19], which consists of 400 English words spoken by a native British English female speaker. The set of words consisted of nouns and verbs (e.g. *talk, claim*), some of which were past-tense inflected (e.g. *arrived, jumped*). We assume that the words’ linguistic properties are independent of the acoustic-phonetic properties presently under investigation. We also assume that this sample of recorded speech provides a reasonable representation of naturally occurring phonetic variants of British English, with the caveat that the sampled utterances are restricted to isolated words and a single speaker.

Audio stimuli, which were originally recorded and presented to subjects with a 22.1 kHz sampling rate, were down-sampled to 16 kHz before building models, as the DNN was trained on a 16 kHz audio training set. After the DNN was first trained on the data from BBC TV programs, it was further adapted to fit the characteristics of the speaker and the recording channel of the stimuli data using an extra adaptation stage with 976 isolated words (see [71] for details of the approach). This is to avoid any potential bias to our experimental results caused by the differences between the DNN model training set and the stimuli set, without requiring the collection of a large amount of speech samples in the same setting as the stimuli set to build a DNN model from scratch. There are no overlapping speech samples (words) between the adaptation and stimuli sets. This guarantees that the model RDM obtained using our stimuli set is not over-fitted into the seen data, and guarantees our results and conclusions to be as general as possible.

### Evaluating clustered representations

Davies–Bouldin indices [14] indicate the suitability of category label assignment to cluster highdimensional data, with lower values indicating better suitability (and with 0 the minimum possible value). To compute Davies–Bouldin indices, we recorded the vector of hidden-layer activations elicited by each input time window of the stimuli for each layer in each DNN. There was a high level of correlation between many activation vectors resulting from overlapping adjacent input vectors. To minimise the effect of this, we used average vectors from each hidden layer over each contiguous phonetic segment. For example, in the word “bulb”, the hidden-layer representations associated with each frame corresponding to the acoustic implementation of the first [b] were combined, and separately the representations for the final [b] were combined. Then, to each combined vector, we assigned a label under five separate labeling schemes: closeness features, frontness features, place features, manner features, and phonetic label. For place and manner features, we considered only phones which exhibited a place or manner feature (*i*.*e*. obstruents). For frontness and closeness features, we likewise considered only phones which exhibited frontness or closeness features (*i*.*e*. syllabic vowels). Where a phone had more than one appropriate feature assignment, we used the most appropriate feature. The full assignment of feature labels for phones used in the clustering analysis is given in S1 Fig.

We computed *p*-values for each Davies–Bouldin index calculation using a permutation procedure in which phone labels were randomized after averaging activation vectors for each segment of input (5,000 permutations). *p*-values were computed by randomizing the labels and recomputing Davies–Bouldin indices 5,000 times, building a distribution of Davies–Bouldin indices under the null hypothesis that phone and feature labels did not systematically explain differences in hidden-layer activations. In all cases, the observed Davies–Bouldin index was lower than the minimum value in the null distribution, yielding an estimated *p*-value of exactly 0.0002. Since the precision of this value is limited by the number of permutations performed, we report it as *p <* 0.001. All Davies–Bouldin index values reported were significant at the *p <* 0.001 level.

### Computing model RDMs from incremental machine states

To encapsulate the representational space of each of the DNN’s hidden layer representations through time, we computed model RDMs from the activation of each layer using the following procedure, illustrated in Fig 5. RSA computations were performed in Matlab using the [46] RSA toolbox.

As described previously, the input layer of the DNN had access to 125 ms of audio input at each time step, to estimate the triphone-HMM-state likelihoods. Since we can only compute model RDMs where the DNN has activations for every word in the stimuli set, only the activations corresponding to the frames whose ending time is smaller than 285 ms (the duration of the shortest word) are used in our experiments Since each frame has a 25 ms duration and a 10 ms shift, only the activations of the first 27 frames of each word are reserved to construct our model RDMs (as the frame index *t* is required to satisfy 10 *× t* ⩽ 25 285).

For each fixed position of the sliding time window on each pair of our 400 stimulus words, we obtained the pattern of activation over the nodes in a particular layer of the DNN. By computing Pearson’s correlation distance (1 *− r*) between activation pattern for each pair of words, we built a 400 *×* 400 model RDM whose rows and columns were indexed by the stimulus words. Then, by moving the sliding time window in 10 ms increments and recomputing model RDM frames in this way, we produced a series of model RDMs which varied throughout the first 260 ms of the stimuli. We repeated this procedure for each hidden layer L2–L7, as well as the input and output layers FBK and TRI, producing in total eight series of model RDMs, or 216 individual model RDM frames. When building a model RDM frame from the input layer FBK, we used only the 40 log-mel filterbank values within the central 25 ms window (and did not include the first derivatives or overlapping context windows).

## Brain mapping

### EMEG data collection

Sixteen right-handed native speakers of British English (six male, aged 19–35 years, self-reported normal hearing) participated in the study. For each participant, recordings of 400 English words, as spoken by a female native British English speaker, were presented binaurally. Each word was repeated once. The study was approved by the Peterborough and Fenland Ethical Committee (UK). Continuous MEG data were recorded using a 306 channels VectorView system (Elektra-Neuromag, Helsinki, Finland). EEG was recorded simultaneously from 70 Ag-AgCl electrodes placed within an elastic cap (EASYCAP GmbH, Herrsching-Breitbrunn, Germany) according to the extended 10/20 system and using a nose electrode as the recording reference. All data were sampled at 1 kHz with a band-pass filter from 0.03 Hz to 330 Hz. Details of the EMEG procedure can be found in [19].

### EMEG source estimation

In order to track the cortical locations of brain–model correspondence, we estimated the location of cortical sources using the anatomically constrained MNE [23] with identical parameters to those used in [19, 56, 63]. MR structural images for each participant were obtained using a GRAPPA 3D MPRAGE sequence (TR = 2250 ms; TE = 2.99 ms; flip-angle = 9 deg; acceleration factor = 2) on a 3 T Trio (Siemens, Erlangen, Germany) with 1 mm isotropic voxels. From the MRI data, a representation of each participant’s cerebral cortex was constructed using FreeSurfer software (https://surfer.nmr.mgh.harvard.edu/). The forward model was calculated with a three-layer boundary element model using the outer surface of the scalp as well as the outer and inner surfaces of the skull identified in the anatomical MRI. This combination of MRI, MEG, and EEG data provides better source localization than MEG or EEG alone [42].

The constructed cortical surface was decimated to yield approximately 12,000 vertices that were used as the locations of the dipoles. This was further restricted to the bilateral superior temporal mask as discussed previously. After applying the bilateral region of interest mask, 661 vertices remained in the left hemisphere and 613 in the right. To perform group analysis, the cortical surfaces of individual subjects were inflated and aligned using a spherical morphing technique implemented by MNE [20]. Sensitivity to neural sources was improved by calculating a noise covariance matrix based on the 100 ms pre-stimulus period. The activations at each location of the cortical surface were estimated over 1 ms windows.

This source-reconstructed representation of the electrophysiological activity of the brain as the listeners heard the target set of 400 words was used to compute brain RDMs.

### Computing brain RDMs in a spatiotemporal searchlight

To match the similarity structures computed from each layer of the DNN to those found in human participants, in the ssRSA procedure, RDMs were calculated from the EMEG data contained within a regular spatial searchlight patch and fixed-width sliding temporal window. We used a patch of vertices of radius 20 mm, and a 25 ms sliding window to match the 25 ms frames used in ASR. The searchlight patch was moved to centre on each vertex in the masked source mesh, while the sliding window is moved throughout the epoch in fixed time-steps of 10 ms. From within each searchlight patch, we extracted the spatiotemporal response pattern from each subject’s EMEG data. We computed word-by-word RDMs using Pearson’s correlation distance (1 *− r*) on the resulting response vectors. These RDMs were averaged across subjects, resulting in one brain RDM for each within-mask vertex. Our 25 ms ssRSA sliding window moved in increments of 10 ms throughout an EMEG epoch of [0, 540] ms, giving us a series of RDMs at each vertex for sliding windows [*t, t* + 25] ms for each value of *t* = 0, 10, …, 510. In total, this resulted in a total of 66,300 brain RDM frames. By using the ssRSA framework, we make this vast number of comparisons tractable by systematising the comparison.

### Systematic brain–model RDM comparisons

The model RDMs computed from the DNN layer activations describe the changing representational dissimilarity space of each layer throughout the duration of the stimulus words. We can think of this as a dynamic model timeline for each layer; a collection of RDMs indexed by time throughout the stimulus. Similarly, the brain data-derived RDMs computed from brain recordings describe the changing representational dissimilarity space of the brain responses at each searchlight location throughout the epoch, which we can think of as a dynamic data timeline. It takes non-zero time for vibrations at the eardrum to elicit responses in auditory cortex (Fig 6A). Therefore, it does not make sense to only compare the DNN RDM from a given time window to the precisely corresponding brain RDM for the same window of stimulus: to do so would be to hypothesize instantaneous auditory processing in auditory nerves and in the brain.

**Figure 6:**
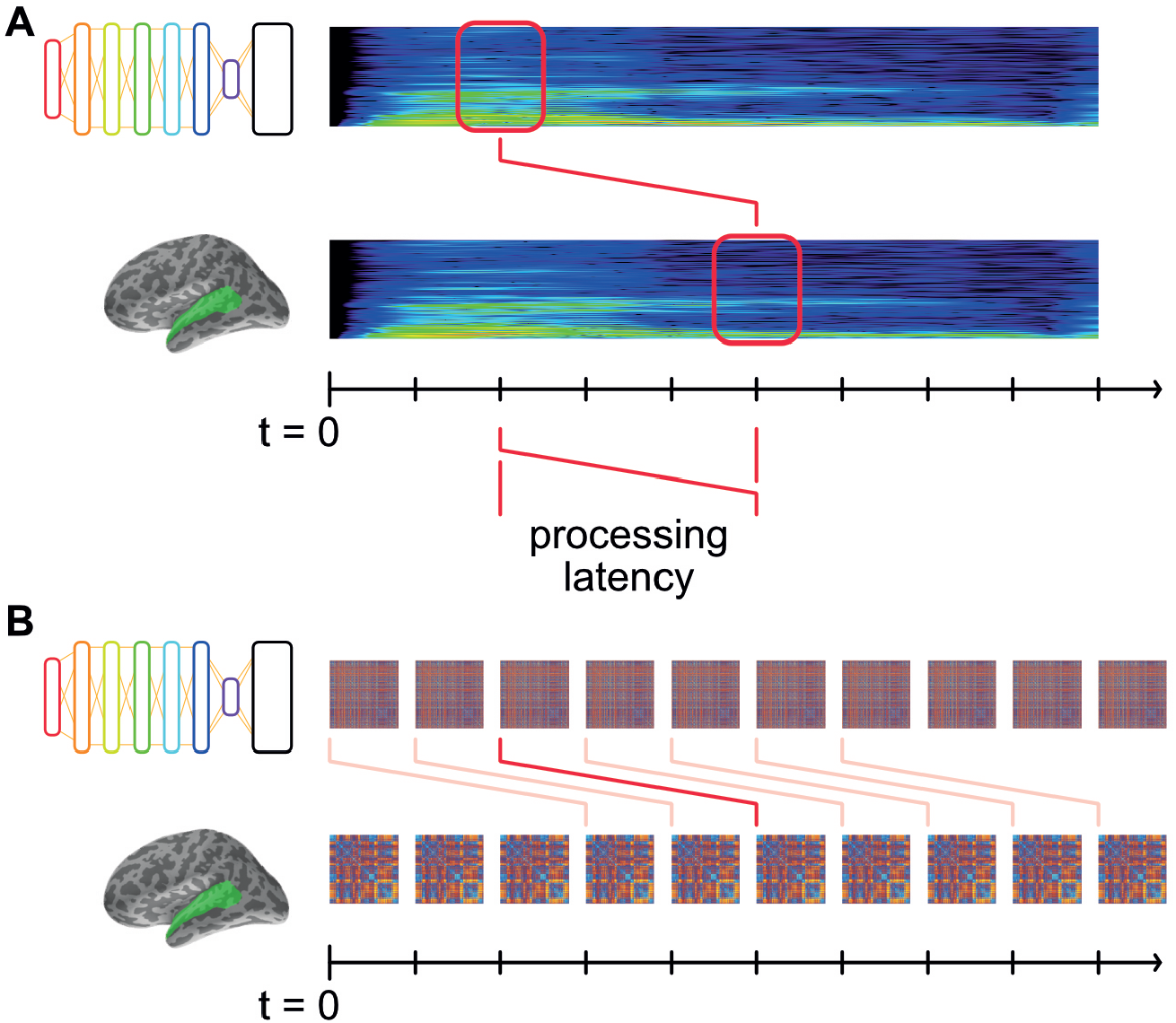
Matching model and data RDMs at systematic latencies. **(A)** Both DNN and brain representations change throughout the time-course of the stimulus, and are aligned to the start of the stimulus at *t* = 0. Some amount of time (“processing latency”) elapses between the sound reaching the participants’ eardrums and the elicited response in auditory cortex. Thus the brain representations recorded at time *t* were elicited by the stimulus earlier in time. **(B)** For a given hypothesized processing latency, we RDMs from DNN layers and brain recordings are matched up, and an overall level of fit is computed. This modelled latency is systematically varied, the resultant level of fit thereby indicating how well the DNN’s representation matches the brain’s at that latency.

Instead, we offset the brain RDM’s timeline by a fixed latency, *k* ms (Fig 6B). Then, matching corresponding DNN and brain RDMs at latency *k* tests the hypothesis that the DNN’s representations explain those in auditory cortex *k* ms later. By systematically varying *k*, we are able to find the time at which the brain’s representations are best explained by those in the DNN layers.

Thus, for each such potential processing latency, we obtain a spatial map describing the degree to which a DNN layer explains the brain’s representations at that latency (i.e. mean Spearman’s rank correlation coefficient between DNN and brain RDMs at that latency). Varying the latency then adds a temporal dimension to the maps of fit.

This process is repeated for each subject, and data combined by a *t*-test of the *ρ* values across subjects at each vertex within the mask and each latency. This resulted in one spatiotemporal *t*-map for each layer of the DNN. For this analysis, we used latencies ranging from 0 ms to 250 ms, in 10 ms increments.

We applied threshold-free cluster enhancement (TFCE: [54]) to the *t*-maps from each layer of the DNN. TFCE is an image-enhancement technique which enables the use of cluster-sensitive statistical methods without the requirement to make an arbitrary choice of initial cluster-forming threshold and is used as the standard statistical method by the FSL software package [26]. All *t*-maps presented for the remainder of this paper have TFCE applied (see S4 Appendix for details).

### Group statistics and correction for multiple comparisons

To assess the statistical significance of the *t*-maps, we converted the *t*-values to *p* values using a random-effects randomisation method over subjects, under which *p*-values are corrected for multiple spatiotemporal comparisons [45, 54, 55]. In the random-effects test, a null-distribution of *t*-values is simulated under the null hypothesis that Spearman’s rank correlation values *ρ* are symmetrically distributed about 0 (*i*.*e*. no effect). By randomly flipping the sign of each individual subject’s *ρ*-maps before computing the *t*-tests across subjects and applying the TFCE transformation, we simulate *t*-maps under the null hypothesis that experimental conditions are 536 not differentially represented in EMEG responses. From each such simulated map, we record the map-maximum *t*-value, and collect these into a null distribution over all permutations. For this 538 analysis we repeated the randomisation 1000 times, and collected separate null distributions for each hemisphere. To assess the statistical significance of a true *t*-value, we see in which quantile it lies in the simulated null distribution of map-maximum randomisation *t*-values.

We performed this procedure separately for the models derived from each layer of the DNN, allowing us to obtain *t*-maps which could be easily thresholded at a fixed, corrected *p*-value.

### Improving DNN design

From the maximum cluster extents of the DNN layers shown in Figure 2, the activations of the DNN acoustic model significantly correspond to the activity in the left-hemisphere of human brain when listening to the same speech samples. This suggests that the DNN and human brain rely on similar mechanisms and internal representations for speech recognition.

Human speech recognition still has superior performance and robustness in comparison to even the most advanced ASR systems, so we reasoned that it could be possible to improve the DNN model structure based on the evidence recorded from the brain.

From the maximum cluster extents of layer L5 in Figure 3A and Figure 3B, hidden layer L5 of the DNN model has a much smaller overall fit to both the left and right hemisphere, compared to the other layers. This indicates the possibility that the calculations in DNN layer L5 are less important for recognising the speech accurately since brain does not appear to use such representations in the recognition process. On the other hand, although a bottleneck layer is positioned at L7, its strong correspondence to the brain reveals the importance of the calculations performed in that layer. Thus, it is natural to assume that more parameters and calculations in important layers can improve speech recognition performance, while fewer calculations can reduce the complexity of the model DNN structure without sacrificing the performance too much.

We verified this by building new DNN models with the bottleneck layer in different positions, controlling the number of parameters by scaling the sizes of the hidden layers in the new DNNs. All the training and test procedures are kept to be the same as previously described. The details of the new DNN structures are shown in Figure 3C.

We tested the derived DNN models with different bottleneck layer positions using two tasks: general large-vocabulary continuous speech recognition with recordings from BBC TV programs, and in-domain isolated-word recognition using the stimuli set. The MGB Dev set was derived as a subset of the official development set of the MGB speech recognition challenge [4], which includes 5.5 hours of speech. Since the MGB testing set involves sufficient samples (8,713 utterances and 1.98M frames) from 285 speakers and 12 shows with diversified genres, and the related WER results are reliable metrics to evaluate the general performance of the DNN models for speech recognition. In contrast, the WERs on the stimuli set are much more noisier since it only consists of 400 isolated words from a single female speaker. However, the stimuli set WERs are still important metrics since the same 400 words are used to build the RDMs used in the key experiments. These results are presented in Table 1 and Fig 4C.

## Supporting information

**S1 Fig.**
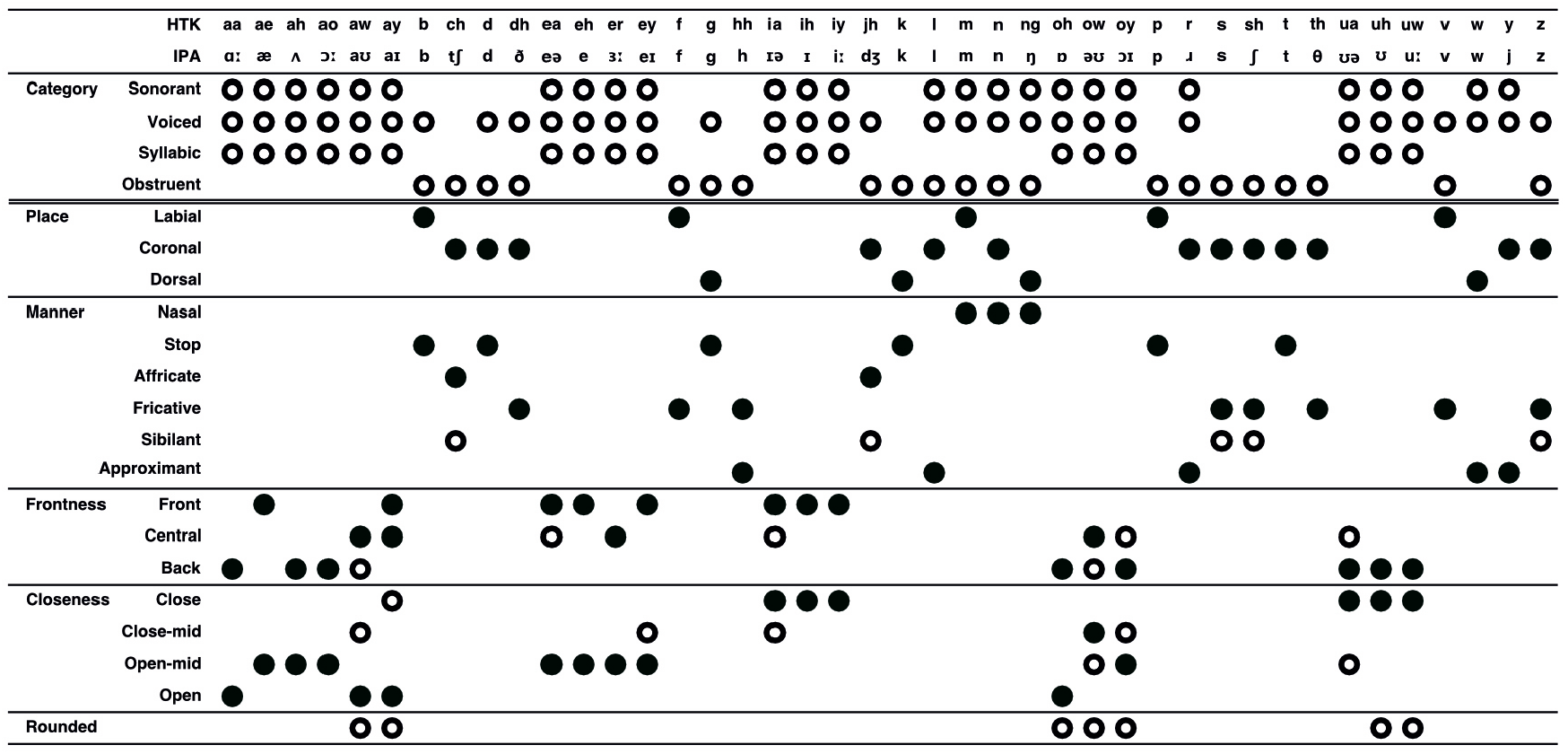
Phone–feature matrix: Assignment of features to phones. Empty circles indicate presence of the feature for a phone. Where a phone has more than one feature for a given category, full circles indicate the dominant feature, used in clustering analysis.

**S2 Table.** Clusters of fit for DNN-BN_7_ in EMEG study. Latencies for left- and right-hemisphere clusters (*p <* 0.01) for each hidden-layer model.

Left hemisphere
| DNN layer model | Cluster latency (ms) |  |  | Peak extent (vertices) |
| --- | --- | --- | --- | --- |
|  | Start | Max | End |  |
| FBK (early cluster) | 0 | 20 | 70 | 93 |
| FBK (late cluster) | 150 | 180 | 200 | 100 |
| L2 | 0 | 160 | 230 | 270 |
| L3 | 0 | 170 | 250 | 237 |
| L4 | 140 | 170 | 200 | 43 |
| L5 |  |  | (n.s.) |  |
| L6 | 20 | 180 | 230 | 151 |
| L7 | 40 | 170 | 230 | 129 |
| TRI |  |  | (n.s.) |  |

Right hemisphere
| DNN layer model | Cluster latency (ms) |  |  | Peak extent (vertices) |
| --- | --- | --- | --- | --- |
|  | Start | Max | End |  |
| FBK | 0 | 170 | 120 | 186 |
| L2 | 0 | 70 | 110 | 172 |
| L3 | 0 | 0 | 110 | 341 |
| L4 | 0 | 0 | 110 | 411 |
| L5 | 0 | 50 | 70 | 20 |
| L6 | 0 | 50 | 120 | 264 |
| L7 | 0 | 40 | 120 | 286 |
| TRI |  |  | (n.s.) |  |

**S3 Fig.**
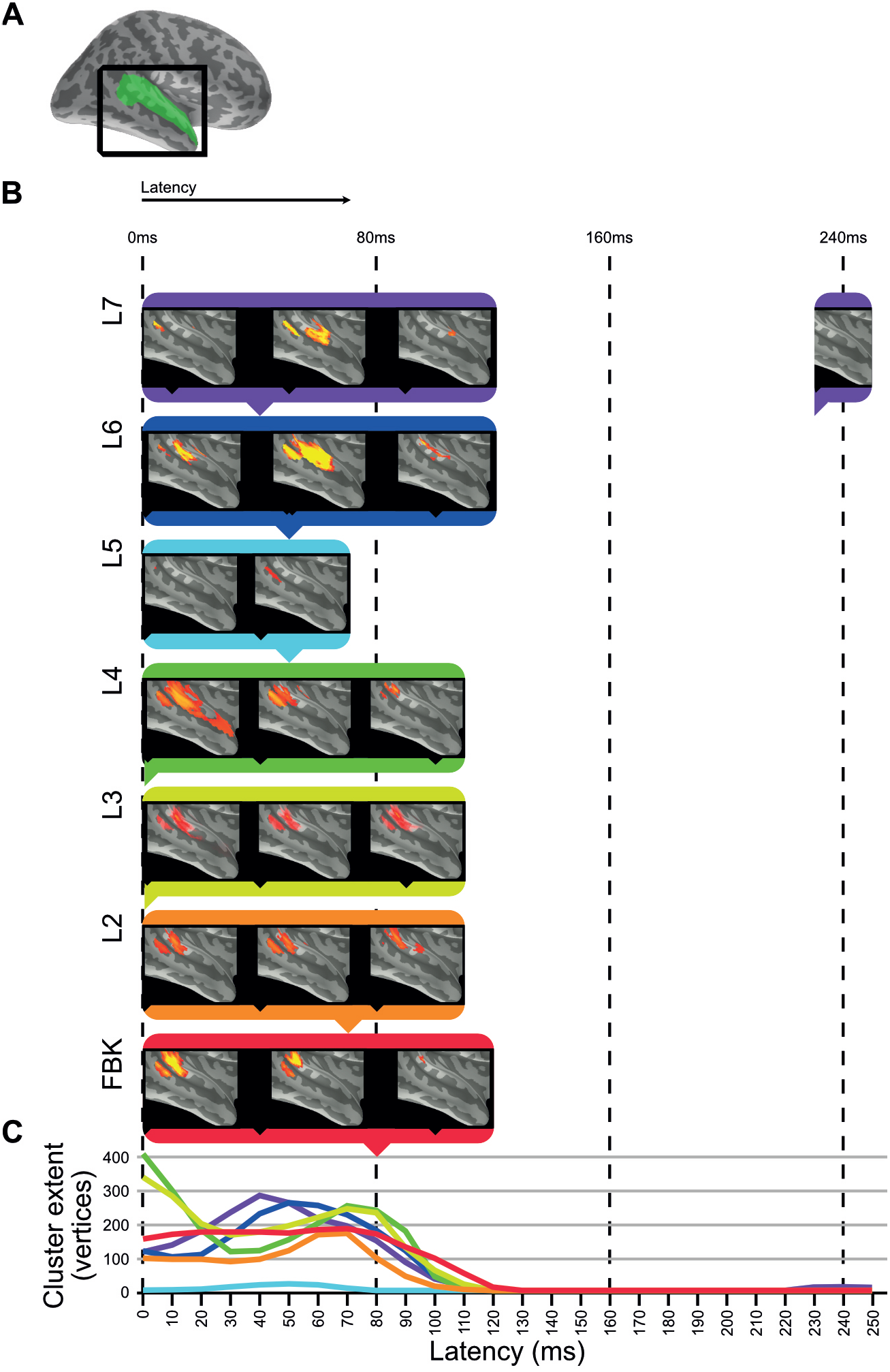
Clusters of significant fit of hidden-layer models to right-hemisphere EMEG data. **(a)** Location of region of interest mask for auditory cortex. **(b)** Maps describing fit of DNN layer models to EMEG data. All maps thresholded at *p <* 0.01 (corrected). **(c)** Line graphs showing the time-courses of cluster extents for each layer which showed significant fit.

**S4 Appendix. Threshold-free cluster enhancement** Threshold-free cluster enhancement (TFCE: [54]) transforms a statistical image in such a way that the value at each point becomes a weighted sum of local supporting clustered signal. Importantly, the shape of isocontours, and hence locations of local maxima, are unchanged by the TFCE transformation. For a *t*-map comprised of values *t*_*v,k*_ for vertices *v* and latencies *k*, the TFCE transformation is given by

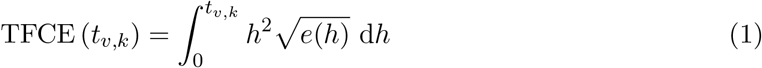

where *e*(*h*) is the cluster extent of the connected component of (*v, k*) at threshold *h*. We approximated (1) with the sum

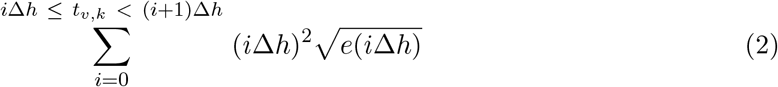

where Δ*h* was set to 0.1. The choice of Δ*h* affects the accuracy of the approximation (2) but should not substantially bias the results.

## Acknowlegements

The authors thank Anastasia Klimovich-Smith, Hun Choi, Lorraine Tyler, Andreas Marouchos and Geoffrey Hinton for thoughtful comments and discussions. RSA computation was done in the RSA toolbox for Matlab [46] using custom EMEG and ssRSA extensions, to which Isma Zulfiqar, Fawad Jamshed and Jana Klímová also contributed. This research was supported financially by a Senior Research Fellowship to LS from Alzheimer’s 601

Research UK (ARUK-SRF2017B-1), an Advanced Investigator grant to WMW from the European Research Council (AdG 230570 NEUROLEX), by MRC Cognition and Brain Sciences Unit (CBSU) funding to WMW (U.1055.04.002.00001.01), and by a European Research Council Advanced Investigator grant under the European Community’s Horizon 2020 Research and Innovation Programme (2014-2020 ERC Grant agreement no 669820) to Lorraine K. Tyler.

## Author contributions

**CW** Conceptualization, formal analysis, software, methodology, writing, editing.

**CZ** Conceptualization, formal analysis, software, methodology, writing, editing, data curation.

**BD** Methodology, editing.

**EF** Data acquisition, data curation, editing.

**AT** Software, data curation, editing.

**XL** Software, methodology, editing.

**PW** Conceptualization, funding acquisition, supervision, editing.

**WMW** Conceptualization, funding acquisition, supervision, editing.

**LS** Conceptualization, software, methodology, funding acquisition, supervision, writing, editing.

## Data availability

The 200 hours Multi-genre Broadcast (MGB) dataset used to train the ASR model was only available to the 2015 MGB-1 Challenge (http://www.mgb-challenge.org/) participants with copyright restrictions from BBC. The 26-dimensional hidden layer representations extracted from the L7 layer of the DNN-BN_7_ model can be found in (http://mi.eng.cam.ac.uk/~cz277/stimuli). Masked, preprocessed human neuroimaging data used for this analysis is available from figshare (https://doi.org/10.6084/m9.figshare.5313484.v1).

## Code availability

The DNN-based ASR system was created using an open-source toolkit, the HTK toolkit version 3.5 (https://htk.eng.cam.ac.uk/). The RSA procedure for this paper was performed using the open-source RSA toolbox (https://github.com/rsagroup/rsatoolbox_matlab), with the addition of specific extensions for ssRSA for EMEG (https://github.com/lisulab/rsatoolbox and https://github.com/lisulab/rsa-dnn-mapping). RDMs were computed from DNN layer representations using publicly available scripts (https://github.com/lisulab/htk-postprocessing).

